# Transcriptome Analysis Reveals Organ-Specific Effects of 2-Deoxyglucose Treatment in Healthy Mice

**DOI:** 10.1101/2023.04.24.537717

**Authors:** Ann E. Wells, John J. Wilson, Sarah E. Heuer, John D. Sears, Jian Wei, Raghav Pandey, Mauro W. Costa, Catherine C. Kaczorowski, Derry C. Roopenian, Chih-Hao Chang, Gregory W. Carter

**Affiliations:** The Jackson Laboratory, Bar Harbor, ME 04469, USA; Tufts University Graduate School of Biomedical Sciences, Boston, MA 02111, USA; Graduate School of Biomedical Sciences and Engineering, University of Maine, Orono, ME 04469, USA

**Keywords:** 2-deoxy-D-glucose, C57BL/6J, metabolism, multi-tissue, transcriptomics

## Abstract

**Objective:** Glycolytic inhibition via 2-deoxy-D-glucose (2DG) has potential therapeutic benefits for a range of diseases, including cancer, epilepsy, systemic lupus erythematosus (SLE), and rheumatoid arthritis (RA), and COVID-19, but the systemic effects of 2DG on gene function across different tissues are unclear.

**Methods:** This study analyzed the transcriptional profiles of nine tissues from C57BL/6J mice treated with 2DG to understand how it modulates pathways systemically. Principal component analysis (PCA), weighted gene co-network analysis (WGCNA), analysis of variance, and pathway analysis were all performed to identify modules altered by 2DG treatment.

**Results:** PCA revealed that samples clustered predominantly by tissue, suggesting that 2DG affects each tissue uniquely. Unsupervised clustering and WGCNA revealed six distinct tissue-specific modules significantly affected by 2DG, each with unique key pathways and genes. 2DG predominantly affected mitochondrial metabolism in the heart, while in the small intestine, it affected immunological pathways.

**Conclusions:** These findings suggest that 2DG has a systemic impact that varies across organs, potentially affecting multiple pathways and functions. The study provides insights into the potential therapeutic benefits of 2DG across different diseases and highlights the importance of understanding its systemic effects for future research and clinical applications.

## 1. Introduction

2-deoxy-D-glucose (2DG) is a glucose analog that has garnered considerable interest in recent years as a potential therapeutic agent for various diseases characterized by abnormal glycolysis, including cancer[1–5], epilepsy[6], systemic lupus erythematosus (SLE)[7], rheumatoid arthritis (RA)[8], and COVID-19 [9–12]. 2DG is taken up into cells by glucose transporters and is converted by hexokinase into 2DG-6-phosphate which cannot be further broken down to yield energy, resulting in a reduction in the rate of glycolysis[13]. Similarly, by competing with mannose, 2DG disrupts the early steps of N-linked glycosylation, ultimately resulting in the misfolding of proteins and the onset of endoplasmic reticulum (ER) stress[14]. Despite an increasing understanding of these cellular mechanisms, there remains a critical need to delineate the specific mechanisms through which 2DG could be used to treat a range of diseases and the organs that would be most affected in this context.

Studies have shown that 2DG can modulate progression or attenuation in multiple disease paradigm. For instance, in anti-tumor effects, 2DG may act through energy restriction to prevent tumor growth while maintaining bodyweight, glucose levels, and immunity[2, 4, 14, 15]. Furthermore, 2DG has been shown to reduce ATP due to its glycolytic properties and protein synthesis through the AMPK/mTORC1 pathway, reducing translation and promoting autophagy[16]. In seizure models, 2DG has been shown to reduce epileptic seizures both acutely and chronically[6]. Additionally, in mice with traumatic brain injuries, cortical slices showed reduced excitatory neurons and prevented epileptiform activity when treated immediately with 2DG[6] . 2DG has also been shown to reduce amyloid precursor protein and amyloid-beta oligomers in an Alzheimer’s mouse model[17], delaying the progression of these critical hallmarks of Alzheimer’s disease. When treating autoimmune diseases, 2DG has attenuated symptoms of SLE, RA, and multiple sclerosis in mice[7, 18]. Additionally, 2DG has been shown to significantly extend the lifespan of SLE-prone mice compared to untreated controls[7]. In these and several other examples, 2DG has shown early promise in both disease prevention and attenuation.

The use of 2DG in cancer therapy has been limited and its therapeutic efficacy has been inconsistent, but it has been shown to be well tolerated with no significant safety concerns raised[5, 13]. Moreover, glycolytic inhibition has been shown to reduce viral replication[11, 19], and 2DG is currently being investigated for its potential to treat severe acute respiratory syndrome coronavirus-2 (SARS-CoV-2)[9–12]. In clinical trials, patients with moderate to severe SARS-CoV-2 infection who were administered oral 2DG showed an improvement in their symptoms and were taken off oxygen supplementation significantly earlier compared to patients treated with standard of care therapies[12]. Although 2DG has shown the potential to ameliorate multiple diseases, a better understanding of 2DG’s systemic effects help to identify potential targets for clinical trials in metabolic disorders.

Much of what is known concerning the effects of 2DG is based on studies of cell lines, selected tissues, whole animal physiology, and disease. The current study, instead, sought to understand the effects of 2DG administered systemically on major organs. Our goal was to understand how 2DG alters baseline metabolism of a genetically uniform population of young, healthy C57BL/6J mice unencumbered by advanced age, disease, or manipulation. Our approach was to analyze the transcriptomes of nine organs (heart, kidney, hippocampus, hypothalamus, prefrontal cortex, skeletal muscle, small intestine, and spleen) from 2DG- and vehicle-treated C57BL/6J mice to assess systemic responses to 2DG treatment (**Fig. 1**). Using a robust statistical filtering strategy, which combined weighted gene co-network analysis (WGCNA), analysis of variance (ANOVA), and correlation, our results show that 2DG altered metabolism, immunity, and transcription in heart, small intestine, hypothalamus, prefrontal cortex, skeletal muscle, and liver through a unique set of genes in each tissue. The small intestine presented an immunomodulatory signature, while both heart and hypothalamus showed reduced mitochondrial metabolism. Few pathways responded to 2DG in skeletal muscle and liver; and all were down-regulated regardless of biological function. Pathways involved in RNA transcription and endoplasmic reticulum (ER) stress in the prefrontal cortex were over and under-expressed in response to 2DG, respectively. Our results provide a comprehensive understanding of the systemic impact of 2DG on glycolysis and N-linked glycosylation and highlight the organ-specific effects through which 2DG acts. Overall, our study lays the foundation for further research on the therapeutic potential of targeting these metabolic pathways.

**Figure 1:**
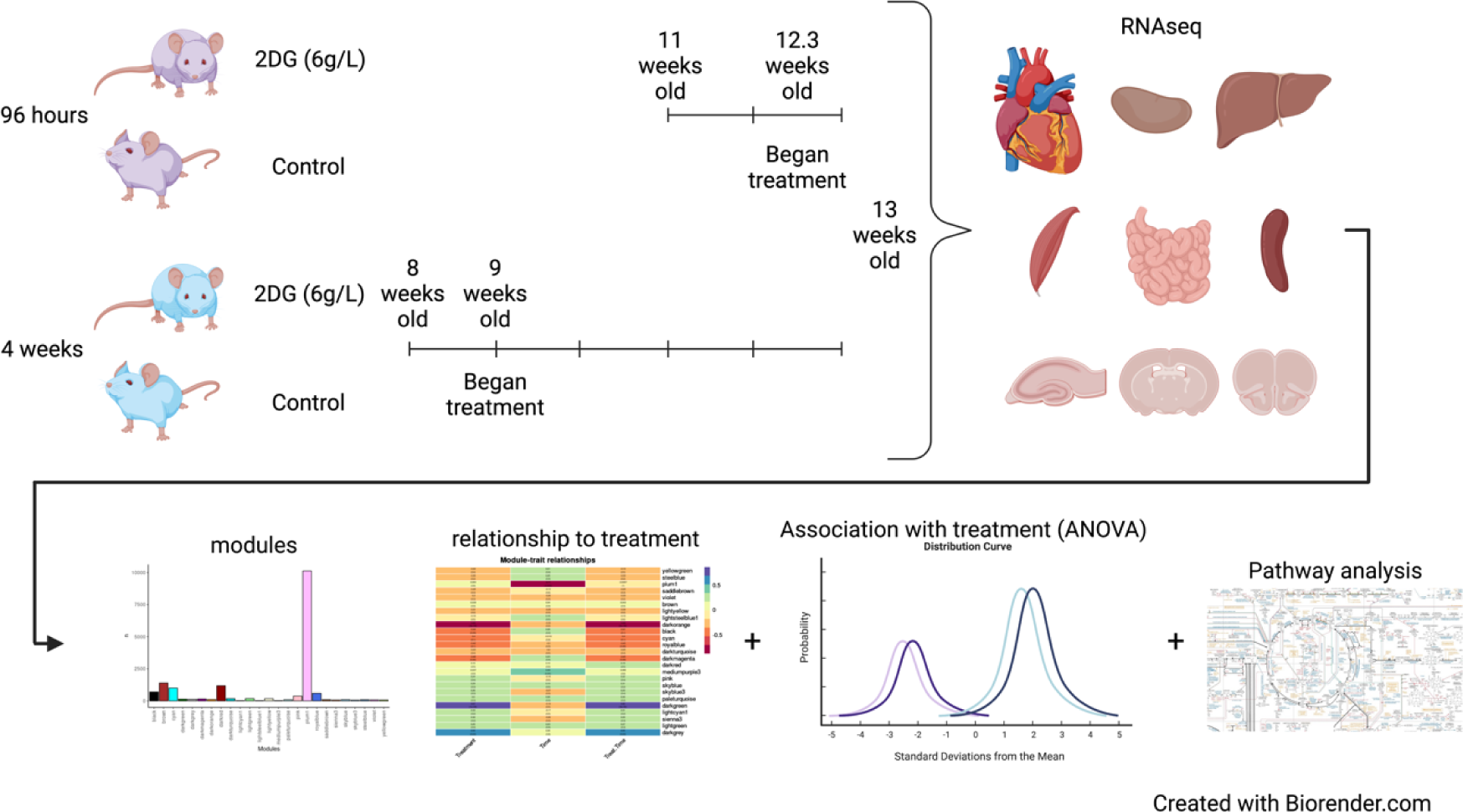
Experimental Design and analysis flow. C57BL/6J mice were treated with 2DG (6g/L) for 96 hours or 4 weeks. Nine tissues were harvested for bulk transcriptomics. Genes identified for each tissue were clustered into modules using WGCNA. Effects of 2DG on each module were determined by assessing each modules relationship with treatment through correlation, ANOVA, and pathway analysis. Created with Biorender.com

## 2. Materials and Methods

### 2.1 Mice, Treatment, and Tissue Isolation

C57BL/6J male mice (Jackson Laboratory, #000664), used in this study were bred and housed at The Jackson Laboratory (Bar Harbor, Maine). Twenty mice at 8 and 20 mice at 11 weeks of age were switched from a 6% (g/g) fat JL Mouse Breeder/Auto 6F (LabDiet**^®^** 5K52) diet to a 10% fat JL Mouse Breeder/Auto (LabDiet**^®^** 5K20) diet when moved into their mouse room, provided with water *ad libitum,* and were housed on a 14-hour light, 10-hour dark cycle in a specific pathogen-free room. 2-Deoxy-D-glucose (2DG, Thermo Fisher Scientific, category number AC111980250, 99% purity) dissolved in drinking water at a concentration of 6 g/L was provided to mice *ad libitum* for either 4-day or 28-day treatment times. As the average mouse consumed ∼ 3.5 ml of water per day, this equates to a dosage of 800 mg/kg/d. The rationale for selecting this dosing regime is that it has proven to be therapeutically effective and safe in treatment of autoimmune disease in several mouse models[18, 20]. After allometric scaling, this dosage approximates an accepted human dosage of ∼65mg/kg/d. The 4-day treatment cohort was provided 2DG-water starting at 12 weeks of age, while the 28-day treatment cohort was provided 2DG-water starting at 9 weeks of age (**Fig. 1**). Control mice received regular drinking water with no additives. All mice, including age-matched control mice, were sacrificed at 13 weeks of age via cervical dislocation. Brains were removed and dissected to isolate the hippocampus, hypothalamus, and pre-frontal cortex. Spleen, liver, heart, kidney, skeletal muscle from left hind-leg and intestinal ileum were also harvested. Heart and intestine were flushed with saline post-dissection to remove blood and feces, respectively. All harvested organs were stored in RNAlater for subsequent RNA-seq analysis. The Jackson Laboratory Institutional Animal Care and Use Committee (IACUC) approved all procedures.

### 2.2 RNA Sequencing

RNA-sequencing (RNA-seq) was performed by Omega Bioservices. Bulk RNA of the 9 excised tissues (heart, hippocampus, hypothalamus, kidney, liver, prefrontal cortex, skeletal muscle, small intestine, and spleen) (4 samples per treatment and time combination) was isolated with the QIAGEN miRNeasy mini extraction kit (QIAGEN) and cDNA was synthesized with the High-Capacity cDNA Reverse Transcription Kit (Applied Biosystems). RNA quality was assessed with a Bioanalyzer 2100 (Agilent Technologies). Poly(A)-selected RNA-seq libraries were generated using the Illumina TruSeq RNA Sample preparation kit v2. RNA-seq was performed in a 150-bp paired-end format with a minimum of 40 million reads per sample on the Illumina HiSeq platform according to the manufacturer’s instructions. RNA-seq reads were filtered and trimmed for quality scores >30 using a custom python script. The filtered reads were aligned to *Mus musculus* GRCm38 using RSEM (v1.2.12) (58) with Bowtie2 (v2.2.0)[21] (command: rsem-calculate-expression -p 12 --phred33-quals --seed-length 25 --forward-prob 0 --time --output-genome-bam -- bowtie2). RSEM calculates expected counts and transcript per million (TPM).

The expected counts from RSEM were used in the Bioconductor edgeR 3.20.9 package[22–24] to determine differentially expressed genes.

### 2.3 Statistical Analysis

All statistical analysis was performed in the language R (3.6.0/4.1.0)[25] equipped with RStudio. The analytical strategy is summarized in Figure 1 and explained in subsequent method sections.

#### 2.3.1 Principal Component Analysis

Principal component analysis (PCA) was performed to identify whether samples clustered based on treatment, time, or treatment-by-time interaction. The PCA algorithm was implemented using the function prcomp() in the package stats[25]. Hotelling’s T^2^ ellipses were used to identify the 95% confidence intervals for each cluster.

#### 2.3.2 Weight Gene Co-network Analysis

To identify potential transcript networks Weighted Gene Co-Expression Network Analysis (WGCNA) was performed using the ‘WGCNA’ R package[26, 27]. Soft-thresholding power was chosen so that the scale-free topology correlation hits as close to 0.9 as possible. The soft thresholding powers chosen for heart, hippocampus, hypothalamus, kidney, liver, prefrontal cortex, skeletal muscle, small intestine, and spleen were 5, 13, 9, 3, 3, 15, 13, and 20, respectively. The adjacency matrix was created using type = “unsigned”. Dynamic tree cutting was used to cluster metabolites and generate modules with a minimum of 15 genes in each module. Networks were identified using all genes quantified by RNA-seq without replacement. Module colors were assigned arbitrarily and have no bearing on functional annotation.

#### 2.3.3 Functional Pathway Analysis

Overrepresentation pathway analysis was performed using gProfiler2[28]. KEGG and reactome databases were used as the references to compare genes in each module identified in WGCNA analysis, using Ensembl 104 and Ensembl Genomes 51 databases. To determine statistical significance of each pathway identified, Fisher’s exact test was used. P-values were adjusted using the Benjamini-Hochberg procedure to reduce type 1 error (FDR < 0.05).

#### 2.3.4 Analysis of Variance using Aligned Rank Transformation

Since the data was not normally distributed aligned rank transformation was performed using the ARTool package[29, 30] to identify significance for using the linear model:

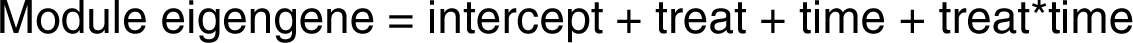

to determine the effects of treatment, time, or treatment-by-time interaction on each module eigengene.

Since the change in diet composition and time points overlap exactly, neither were individually assessed in this study but they were evaluated in the ANOVA models—at the module, gene, and Gene Set Variation Analysis (GSVA) level—for interaction to identify their combined influence on treatment. Only modules, GSVA, and genes significant for treatment but not time or treatment dependent on time were discussed.

#### 2.3.5 Mediation Analysis

The R package mediation[31] was used to perform mediation analysis. Eigengenes for overrepresented pathways were created using GSVA[32] (explained in section 2.6) and assessed across tissues to determine if pathways were acting as mediators on the response to treatment using two models; the mediator model:

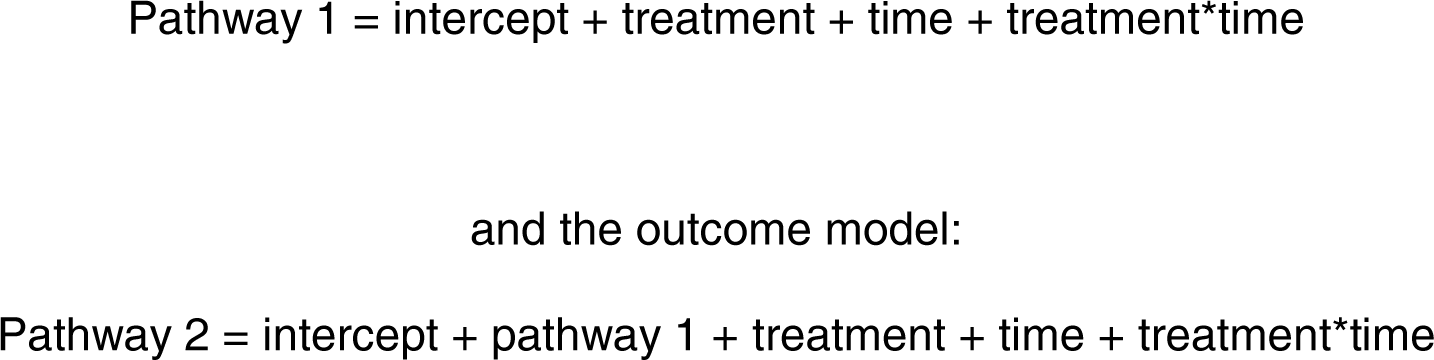

The models were then entered into the function mediate() and simulated 1000 times. Confidence intervals and p-values were created using the Quasi-Bayesian Monte Carlo method.

### 2.4 Modules Identified Using Three Filtering Criteria

To identify modules related to treatment for each tissue each module had to meet three criteria: 1.) module was significantly correlated with treatment, 2.) module identified significantly overrepresented pathways, and 3.) module was uniquely significant for treatment only, according to ANOVA. Correlation was used to directly describe the relationship between each module and treatment. ANOVA was used to assess the difference between main effects and their interactions. This filtering strategy was chosen to identify modules that had the most robust response to 2DG treatment. Modules that fit these criteria were further assessed for genes that may be central to the networks identified.

### 2.5 Gene Set Variation Analysis

GSVA was performed, using the GSVA package[32], to identify enrichment of significantly overrepresented pathways for each tissue module (see data resource for comprehensive description of all modules, section 2.6). To identify enriched pathways a reference set of pathways is needed. To create the reference set of pathways, all ensembl genes listed for *Mus musculus* were compared against the KEGG and reactome databases in gProfiler2 using the gost() function and the ensembl 104 and ensembl genomes 51 databases. This identified 2,030 pathways, which were used as the pathway gene set. The set of genes in each module was then assessed against the pathway gene set to identify significantly overrepresented pathways. To calculate overrepresentation, we used the Fisher’s exact test and corrected for multiple comparisons using the Benjamini-Hochberg procedure (FDR < 0.05). To summarize the gene expression in each pathway, the method plage was used, which uses the coefficients of the first right-singular vector from the singular value decomposition. Genes within these overrepresented pathways were then summarized into a singular eigengene representing their expression for each sample. The pathway eigengenes for each significantly overrepresented pathway were then tested using ANOVA:

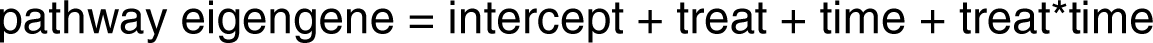

to identify resident pathways significantly altered by treatment. This identified which significantly overrepresented pathways were also significantly enriched.

### 2.6 Data Resource Development for Reproducible and Transparent Data

The data resource was developed using R 4.1.0, rmarkdown, and bookdown[33]. Static html files were output using knitr and collated using a yaml file that controls the navigation bar on the website and the general theme and layout, including footer information, and links in the navigation bar to the shiny app, git repository, figshare, and contact information. The theme used for the website was cerulean and type was set to inverse to change the color to dark blue. The table of contents for each page was set to float. The website is publicly hosted at: https://storage.googleapis.com/bl6_2dg_rnaseq/index.html.

## 3. Results

### 3.1 Molecular changes are induced in a tissue-specific manner by 2-DG treatment

The impact of 2DG on transcriptomic profiles of nine different tissues was assessed to identify genes that were differentially expressed for treatment only. The number of differentially expressed genes varied across the tissues, with a minimum of 230 genes in spleen and a maximum of 1152 genes in hippocampus. Interestingly, only three tissues (muscle (1), small intestine (119), and prefrontal cortex (3)) exhibited overrepresented pathways, indicating that the effect of 2DG on biological pathways may be more significant in these tissues than in the other six tissues examined. Assessing 2DG’s effect at a granular level, however, do not account for potential gene interactions. These findings suggest that assessing the correlation structure of genes within each tissue may provide a better biological understanding of 2DG’s impact at the tissue level.

To identify the most robust signature in each tissue and minimize noise, a novel filtering strategy was devised (**Fig. 1**). Principal Component Analysis showed that samples clustered by tissue based on their transcriptomic profiles (**Fig. 2**). We used WGCNA to group genes into modules that share highly correlated expression patterns. Across tissues, the number of modules into which genes were clustered ranged from 25 to 46 and contained anywhere from 22 to 10,127 genes within each module. Identifying modules that fit the three criteria in the filtering strategy (significant for treatment, through correlation and ANOVA, and contained significantly overrepresented pathways (p < 0.05, FDR < 0.05)), we narrowed our focus to six tissues — heart, hypothalamus, liver, skeletal muscle, prefrontal cortex, and small intestine — and identified one module for each of these tissues. This approach allowed us to elucidate the most robust changes in gene expression in response to 2DG treatment in each tissue and highlight the biological pathways that are most affected. More information about the analysis, including the number of modules and genes per tissue, can be found in the Supplementary Information or on our website (https://storage.googleapis.com/bl6_2dg_rnaseq/index.html).

**Figure 2:**
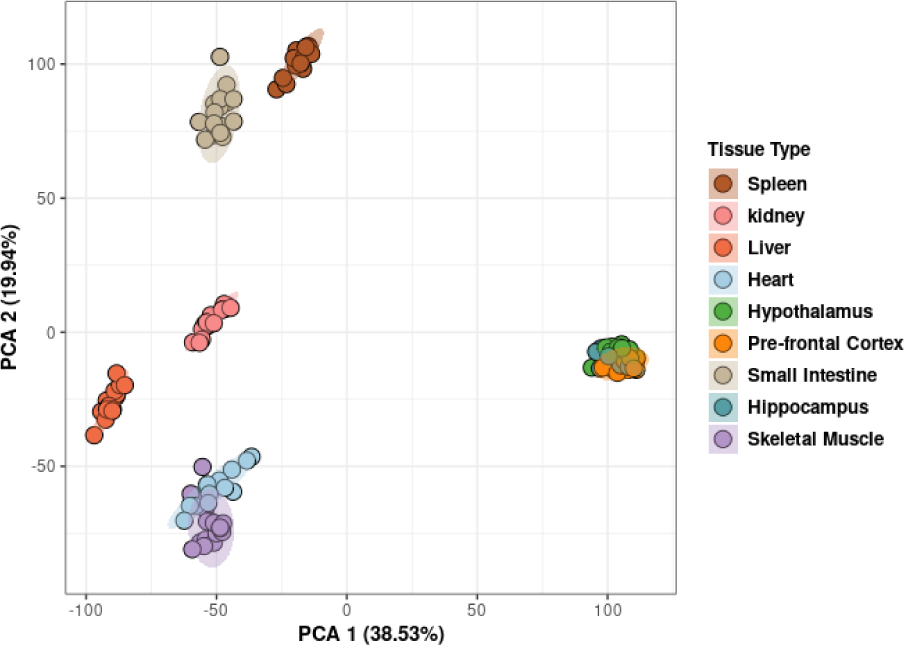
Principal Component Analysis demonstrated clustering of samples based on tissue type. Hotelling’s T^2^ ellipses were calculated, and the clustering patterns of all tissue were contained within the 95% confidence intervals.

### 3.2 2DG reduced expression of metabolic pathways in the heart

The expression of genes in heart muscle was analyzed and found to cluster into 25 distinct modules (**Supplementary Fig. 1A**), with three of them showing significant correlation with 2DG treatment (**Supplementary Fig. 1B**). Further analysis using ANOVA revealed that only the darkgreen module was uniquely significant for the treatment (p < 0.001). This module contained 138 genes predominantly related to metabolism (**Supplementary Fig. 1C**), which summary eigengenes revealed were decreased in expression in 2DG-treated mice compared to control mice (**Supplementary Fig. 1D, E**). The darkgreen module was found to contain ten pathways related to mitochondrial metabolism and the breakdown of branched-chain amino acids (BCAA) (**Table 1**), which were significantly overrepresented (p < 0.05). Comparison of genes within each pathway using a Jaccard similarity index revealed that each pathway consisted of a unique set of genes (**Supplementary Fig. 2**). To assess pathway enrichment, gene set variation analysis (GSVA) was performed. Of the ten significantly overrepresented pathways, seven showed significant reduction in eigengene expression for the treatment group only (p < 0.05, **Supplementary Table 1**), referred to as high confidence pathways from this point forward.

**Table 1:** Metabolic pathways are overrepresented in the darkgreen heart module

| Pathways | GSVA Significance for Treatment<br>(ANOVA p-value)* | GSVA Expression<br>Direction (2DG<br>compared to Control) |
| --- | --- | --- |
| Metabolism | **0.0001** | Down |
| Mitochondrial translation | 0.133 | Down |
| Mitochondrial translation<br>elongation | 0.133 | Down |
| Mitochondrial translation<br>termination | 0.133 | Down |
| Propanoate metabolism | **0.033** | Down |
| Pyruvate metabolism | **0.033** | Down |
| Pyruvate metabolism and Citric Acid (TCA) cycle | **0.004** | Down |
| Regulation of pyruvate dehydrogenase (PDH) complex | **0.001** | Down |
| The citric acid (TCA) cycle and respiratory electron transport | **0.001** | Down |
| Valine, leucine, and isoleucine degradation | **0.002** | Up |

\* Significant p-values in bold

BCAA degradation was found to be inversely correlated with changes in mitochondrial metabolism, consistent with previous findings of mitochondrial dysfunction leading to BCAA accumulation[34]. Failure in both mitochondrial metabolism and BCAA degradation have been linked to increase cardiovascular disease[35], and reduced BCAA degradation has been identified as a signature in heart failure[36]. The inverse relationship of BCAA degradation and mitochondrial metabolism identified in this study suggests that 2DG’s differential effects on heart metabolism may provide protection against heart failure.

### 3.3 2DG suppressed expression of immunological pathways in the small intestine

The genes expressed in the small intestine were clustered into 46 modules (**Fig. 3A**). Three of these modules were found to be significantly correlated for treatment (p < 0.05) and identified significantly overrepresented pathways (p < 0.05), with the green4 module being uniquely significant for treatment, according to ANOVA (p = 0.04, **Fig. 3B**). This module contained 467 genes (**Fig. 3E**) predominantly related to the immune system and summary eigengenes revealed decreased expression in mice treated with 2DG compared to control mice (**Fig. 3C, D**). Overrepresentation analysis identified 69 significant pathways (p < 0.05, **Supplementary Table 2**), containing a unique set of genes in each pathway according to the Jaccard similarity index (**Supplementary Fig. 3**). GSVA results showed that 61 out of the 69 pathways were significantly enriched for treatment only (p < 0.05, **Supplementary Table 2**) and therefore indicated high confidence. Furthermore, when pathways were functionally annotated it was noted that 39 of the 61 high confidence pathways were related to immunological function and immunology related diseases. Thirteen pathways showed decreased gene expression in mice treated with 2DG, while 26 pathways showed increased expression. The up-regulated pathways were primarily involved in innate immunity and the down-regulated pathways were related to adaptive immunity (**Supplementary Table 2**). CIBERSORT analysis was used to assess immune cell types in the green4 module and found that nine cell types were determined to be significantly different between the green4 module and all the genes identified within the small intestine (referred to as all genes). Four of these cell types (eosinophils, macrophages, memory B cells, and Th1 cells) were significantly increased in the green4 module compared to all genes (P < 0.05). The expression of genes related to eosinophils and macrophages (innate immune cells) increased with 2DG treatment, while genes related to memory B cells and Th1 cells (adaptive immune cells) decreased, but not significantly. The expression of genes related to immature dendritic cells was significantly different between the green4 module and all genes but in a treatment-dependent manner, with decreased expression in the green4 module and mice treated with 2DG compared to all genes and control mice (**Supplementary Table 3**). Collectively, these results indicate that treatment with 2DG leads to changes in the gene expression profile of the small intestine’s immune system characterized by a reduction in adaptive immunity, and potentially compensation for corresponding by an enrichment in innate immune cells.

**Figure 3:**
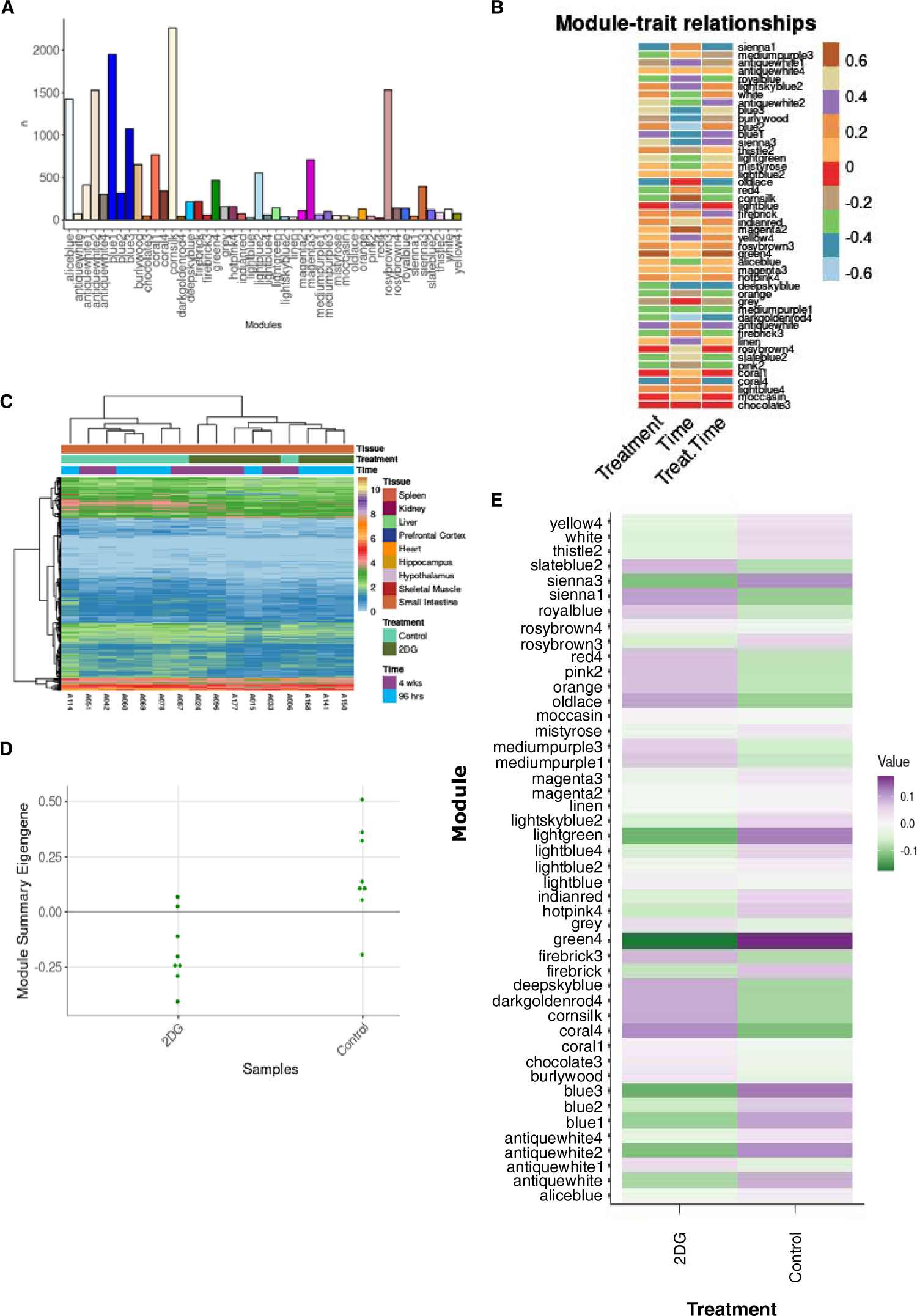
Analysis of modules-treatment association. A) A total of 46 modules were identified in the small intestine, each containing between 22 and 2,259 genes. B) Correlation analysis between time, treatment, and treatment in a time dependent manner revealed three modules significantly correlated with treatment. C) The average eigengene expression of the green4 module shows that expression levels are lower in mice treated with 2DG compared to control mice. D) Summary eigengene expression for each sample in the green4 module confirms lower expression in mice treated with 2DG compared to control mice., and E) Individual genes across samples clustered by treatment except for one control sample, which clustered with 2DG-treated mice.

### 3.4 2DG reduced expression of nicotinate metabolism in the liver

A total of thirty-one modules were identified in liver (**Supplementary Fig. 4A**), but only the darkolivegreen module, consisting of 280 genes that represented diverse functions met all three filtering criteria (**Supplementary Fig. 4B, C**). The summary eigengenes revealed increased expression in 2DG-treated mice compared to the control mice (**Supplementary Fig. 4D, E**). The only significant pathway found was nicotinate metabolism, which was deemed to be of high confidence through GSVA analysis. This pathway was overrepresented (p < 0.05, **Supplementary Table 4**) and was significantly reduced in mice treated with 2DG compared to the control group (p = 0.008). Nicotinate metabolism is important for maintaining levels of nicotinamide adenine dinucleotide (NAD+), an important coenzyme necessary for energy metabolism[37], which is primarily produced in the liver[38]. These findings suggest that 2DG treatment may have an impact on NAD+ levels in the liver and thus on ATP production.

### 3.5 2DG reduced expression of protein metabolism in skeletal muscle

A comprehensive analysis of the skeletal muscle revealed the existence of 40 modules (**Supplementary Fig. 5A**), of which 32 were found to have significantly overrepresented pathways and three were significantly correlated with the treatment (p < 0.05, **Supplementary Fig. 5B**). Further analysis revealed that among these three modules, only the salmon2 module was determined to be unique and significant for treatment through ANOVA analysis (p = 0.017). This module contained 153 genes (**Supplementary Fig. 5C**) and showed an increase in summary eigengenes in mice treated with 2DG compared to the control mice (**Supplementary Fig. 5D, E**). Like the findings in liver, genes in this module were found to be associated with a myriad of functions, but only five pathways related to protein metabolism were significantly overrepresented (p < 0.05, **Supplementary Table 5**) and showed a reduction in expression after 2DG treatment. Further analysis using the Jaccard similarity index and GSVA revealed that the genes involved in each pathway were unique (**Supplementary Fig. 6**), and two of the five pathways were deemed uniquely significant for treatment and therefore high confidence (p < 0.05). As N-linked glycosylation is crucial for protein metabolism[39], the reduction in expression may lead to up-regulation of the unfolded protein response and down-regulation of protein synthesis. Overall, these data suggest that the treatment with 2DG has a significant impact on the expression of genes related to protein metabolism in the skeletal muscle.

### 3.6 2DG increased expression of fatty acid oxidation in the hypothalamus

The genes expressed in the hypothalamus grouped into 25 modules (**Supplementary Fig. 7A**). Four of these modules were found to be correlated with treatment (**Supplementary Fig. 7B**); however, only the bisque4 module was uniquely significant for treatment (p = 0.008). This module contained 289 genes predominantly related to metabolism and cell cycle (**Supplementary Fig. 7C**). The summary eigengenes of this module were decreased in mice treated with 2DG compared to the control mice (**Supplementary Fig. 7D, E**). Nine pathways were significantly overrepresented (p < 0.05), and the Jaccard similarity index showed that the genes identified in each pathway were unique (**Supplementary Fig. 8**). Functional annotation revealed that these nine pathways were related to fatty acid oxidation and chromosome health. Two of the overrepresented pathways (mitochondrial fatty acid beta-oxidation and mitochondrial fatty acid beta-oxidation of saturated fatty acids) were uniquely enriched for treatment based on GSVA analysis, supporting their high confidence (p < 0.05, **Supplementary Table 6**). Both pathways were related to fatty acid oxidation, which often increases as a result of reduced glycolysis[40]. Although the other seven pathways were not significantly enriched using GSVA, the eigengenes in the G2/M DNA damage checkpoint pathway were negatively correlated with each of the other eight pathways (**Supplementary Table 6**). The G2/M DNA checkpoint showed significantly reduced expression in mice treated with 2DG (p = 0.02), but also for time (p = 0.036, **Supplementary Table 6**). These results indicate that the hypothalamus increase fatty acid oxidation in response to the blockade of glycolysis and protects DNA through upregulation of pathways related to chromosome maintenance, potentially in response to the increase in unfolded protein response[41]. Overall, these findings suggests that the hypothalamus may use lipids as an alternative fuel source in response to 2DG.

### 3.7 Enhancement of RNA transcription pathways by 2DG in the prefrontal cortex

Thirty-six gene modules were identified in the prefrontal cortex (**Supplementary Fig. 9A**). Of these, five modules showed a correlation with treatment (p < 0.05, **Supplementary Fig. 9B**) and 26 modules contained significantly overrepresented pathways (p < 0.05), with only the bisque2 module being uniquely significant for treatment, according to ANOVA (p = 0.014). The bisque2 module consisted of 718 genes that represented many biological functions (**Supplementary Fig. 9C**). The summary eigengenes were found to be elevated in 2DG-treated mice compared to control mice (**Supplementary Fig. 9D, E**). Functional analysis revealed that 14 pathways were overrepresented in the bisque2 module (p < 0.05), with seven related to RNA transcription and seven to protein metabolism. According to the Jaccard similarity index, the genes in each pathway were found to be unique (**Supplementary Fig. 10**). Among these pathways, 12 were considered high confidence pathways, as they showed significant enrichment for treatment using GSVA (p < 0.05, **Supplementary Table 7**). 2DG increased the overall expression of five high confidence pathways related to RNA transcription compared to control mice. Conversely, the seven high confidence pathways related to protein metabolism showed decreased expression in 2DG-treated mice compared to control mice. Although 2DG decreased expression of the nucleotide excision repair pathways, it was negatively correlated with other pathways that also showed decreased expression in response to 2DG. All pathways related to RNA transcription were negatively correlated with pathways involved in protein metabolism. These results indicate that in the prefrontal cortex protein metabolism and RNA transcription both respond to 2DG. Their differential response to 2DG may protect the prefrontal cortex from the accumulation of unfolded proteins caused by the disruption of N-linked glycosylation.

### 3.8 Tissue-specific responses to 2DG treatment revealed through correlation analysis of gene modules

Tissues act synergistically to maintain the health of the body, but each tissue expresses genes at varying levels. In this study, the expression of the same genes was analyzed across tissues; however, the comparison of the genes contained in each selected tissue module revealed minimal gene overlap (**Fig. 4**). The highest number of genes shared across tissue modules, although not significant (p = 0.117), was 18 between skeletal muscle and prefrontal cortex, while the number of genes identified in each module ranged from 113 to 659. No gene was shared by all tissues. Moreover, we also compared overrepresented pathways across tissues and identified only one shared pathway between small intestine and prefrontal cortex (**Supplementary Fig. 11**). These results indicate that the tissue-specific effects of 2DG are generally unique at both the gene and pathway levels.

**Figure 4:**
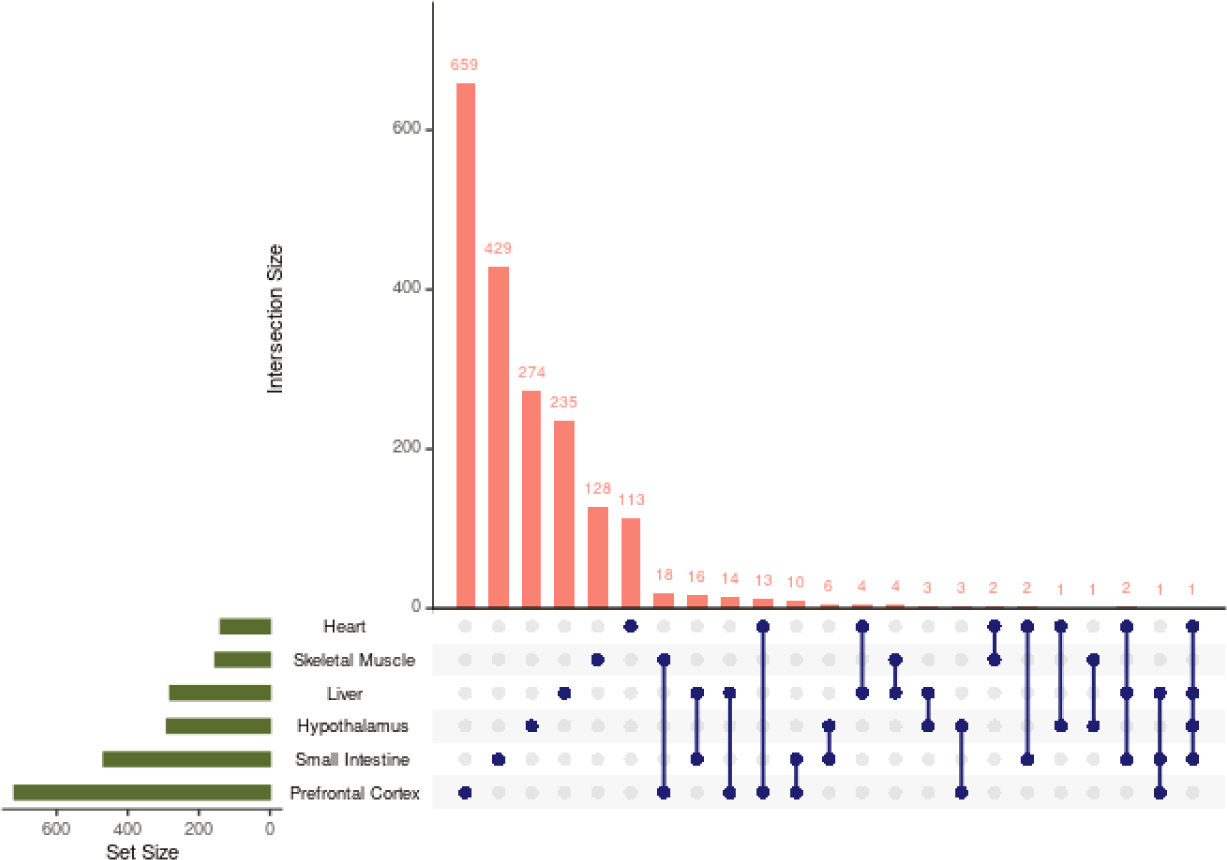
Number of identified genes shared between tissue modules. Each dot represents the tissue or tissues being compared. Very few genes were shared across tissues and no gene was shared across all tissues. Only tissue combinations with one or more shared genes are shown.

We also identified distinct, organ-specific responses to 2DG that can be interpreted as a network of co-occurring transcriptional changes across the organism (**Fig. 5**). To investigate the relationships between tissue modules, we performed correlation analysis, resulting in five significant pairwise relationships out of 21 (p < 0.05, **Fig. 5**). To gain a deeper understanding of these relationships, we used GSVA eigengenes to examine the relationship of each pathway across each tissue module. Positive correlations were found between small intestine and heart tissues (r = .51, p = 0.08). Further assessment at the pathway level revealed a complex relationship between adaptive and innate immunity in the small intestine and BCAA degradation in the heart (**Fig. 5, Supplementary Table 8**). The pathways predominantly related to adaptive immunity were inversely correlated with BCAA degradation, while those predominantly related to innate immunity were positively correlated. Although lymphocytes do not remain in the heart for any length of time, BCAA are necessary for lymphocyte development and clonal expansion[42–45]. To assess any causal relationship between BCAA degradation and pathways in the small intestine we performed causal mediation analysis. BCAA degradation did not mediate any pathway’s response to 2DG in the small intestine. These results indicate that while there is an association between adaptive and innate immunity in the small intestine and BCAA degradation in heart there is not causing the response to 2DG in the small intestine.

**Figure 5:**
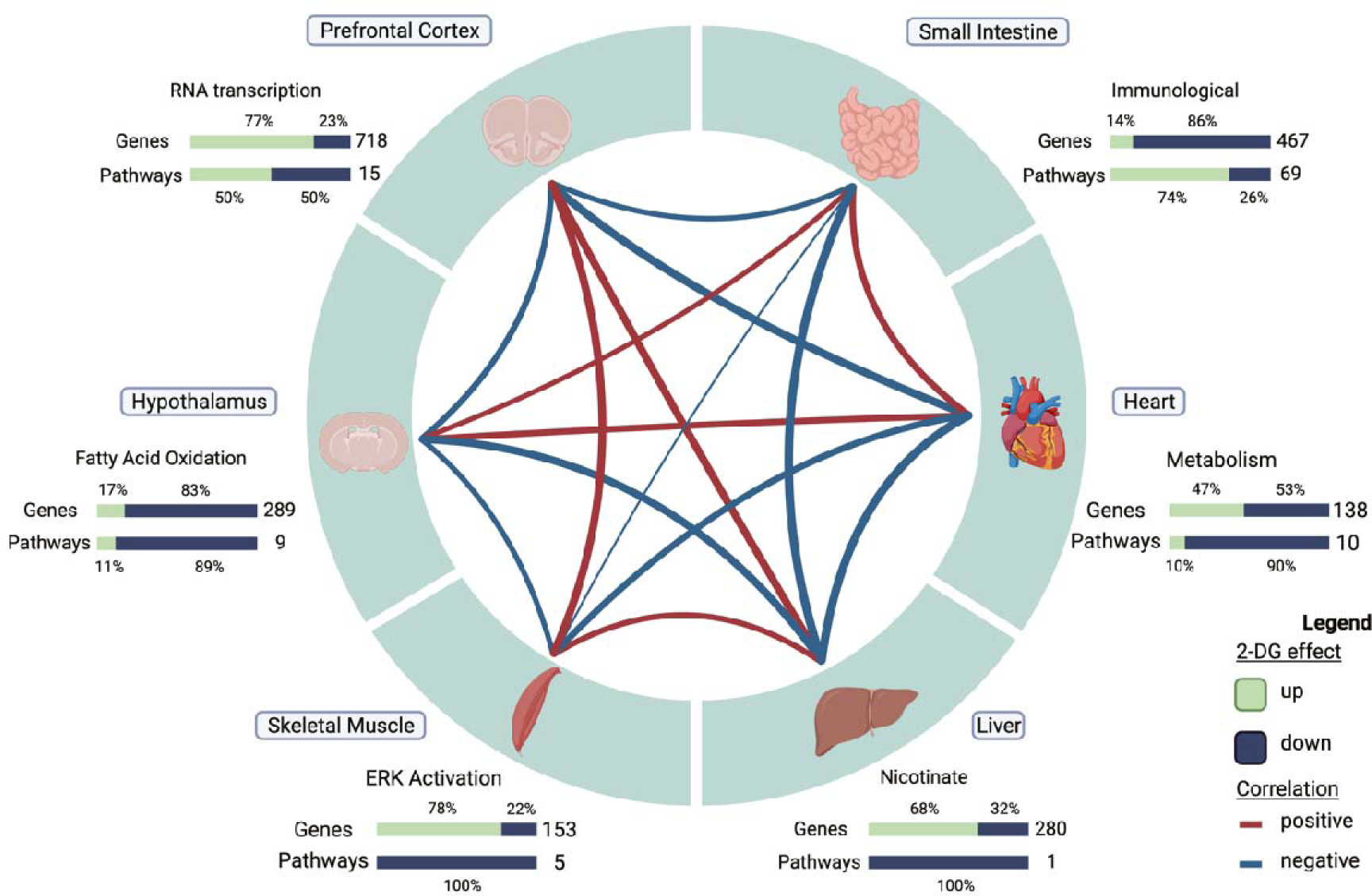
Correlation structure of gene module changes across tissues in 2DG-treated mice. Six tissues, each represented by one high confidence module, demonstrated unique response to 2DG treatment. The expression levels of genes and pathways within each tissue showed varying degrees of over- and under-expression in response to 2DG. Created with Biorender.com.

Tissue modules of the prefrontal cortex and hypothalamus were inversely related (r = -0.48, p = 0.09, **Fig. 6**). The incorporation of 2DG into the oligosaccharide chain inhibits N-linked glycosylation and can lead to the formation of unfolded glycoproteins and induce ER stress[19]. In response to ER stress, the cell cycle is arrested at the G2/M DNA damage checkpoint[46].

**Figure 6:**
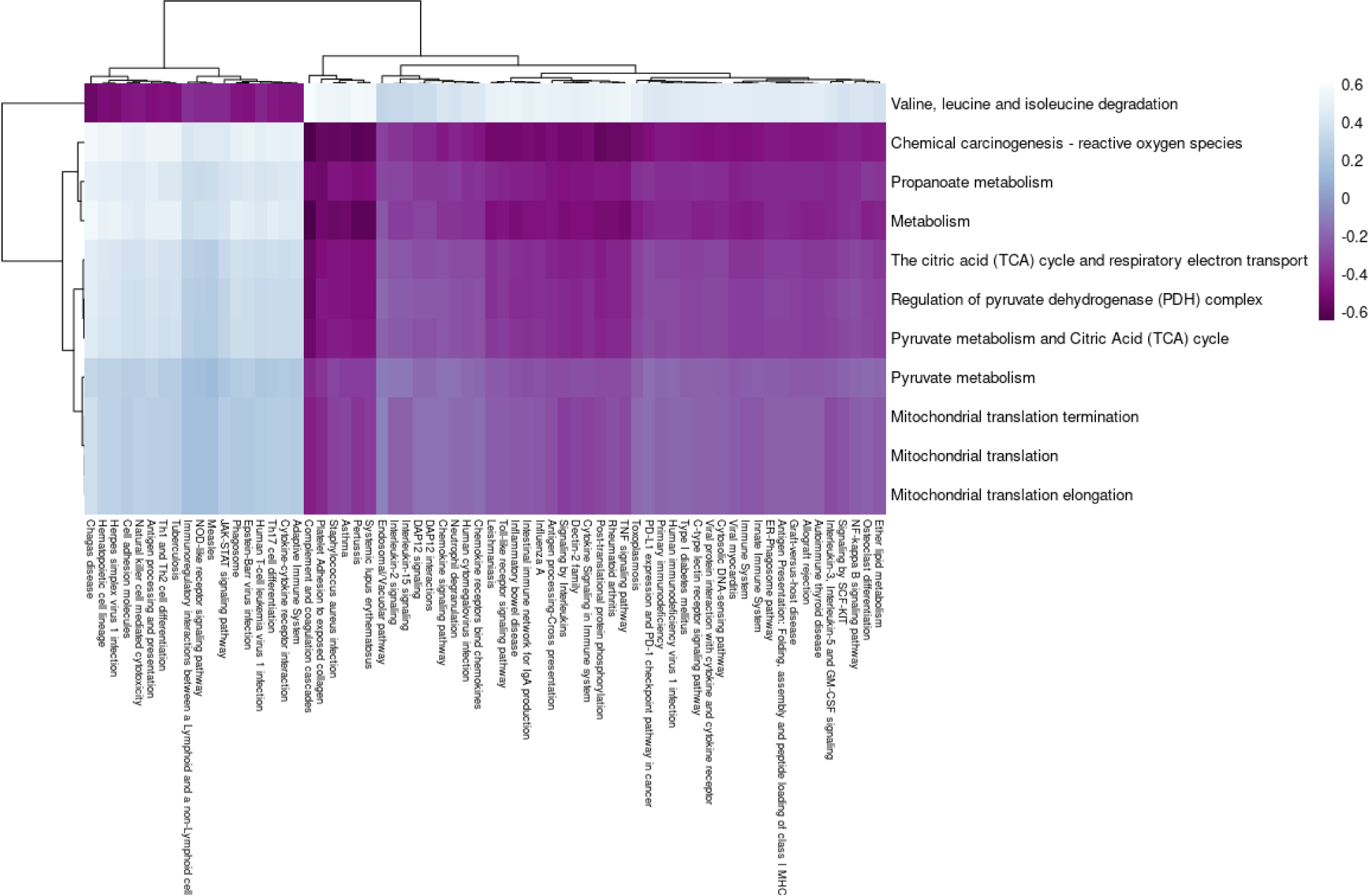
Correlations between pathway eigengenes in the heart (y-axis) and small intestine (x-axis). Branched chain amino acid degradation was negatively correlated with pathways predominantly related to the adaptive immunity and positively correlated with pathways predominantly related to innate immunity in small intestine. All other metabolic pathways in heart were negatively correlated with pathways related to innate immunity and positively correlated with pathways related to adaptive immunity in small intestine.

Our findings showed a positive correlation between the G2/M DNA checkpoint pathway in the hypothalamus and pathways related to ER stress in the prefrontal cortex (**Fig. 7, Supplementary Table 9**). To assess the potential causal relationship between the hypothalamus and prefrontal cortex, we conducted a causal mediation analysis and observed that the G2/M DNA damage checkpoint pathway had a direct effect across all pathways in the prefrontal cortex (p < 0.05) but no causal mediated effects. This indicates that the activation of the G2/M DNA damage checkpoint pathway may have been a secondary response to the ER stress induced by 2DG treatment, rather than the primary cause of the differential response to 2DG. Overall, these results suggest that both the hypothalamus and prefrontal cortex respond to the disruption of N-linked glycosylation by modulating the expression of pathways to counteract ER stress and the accumulation of unfolded proteins.

**Figure 7:**
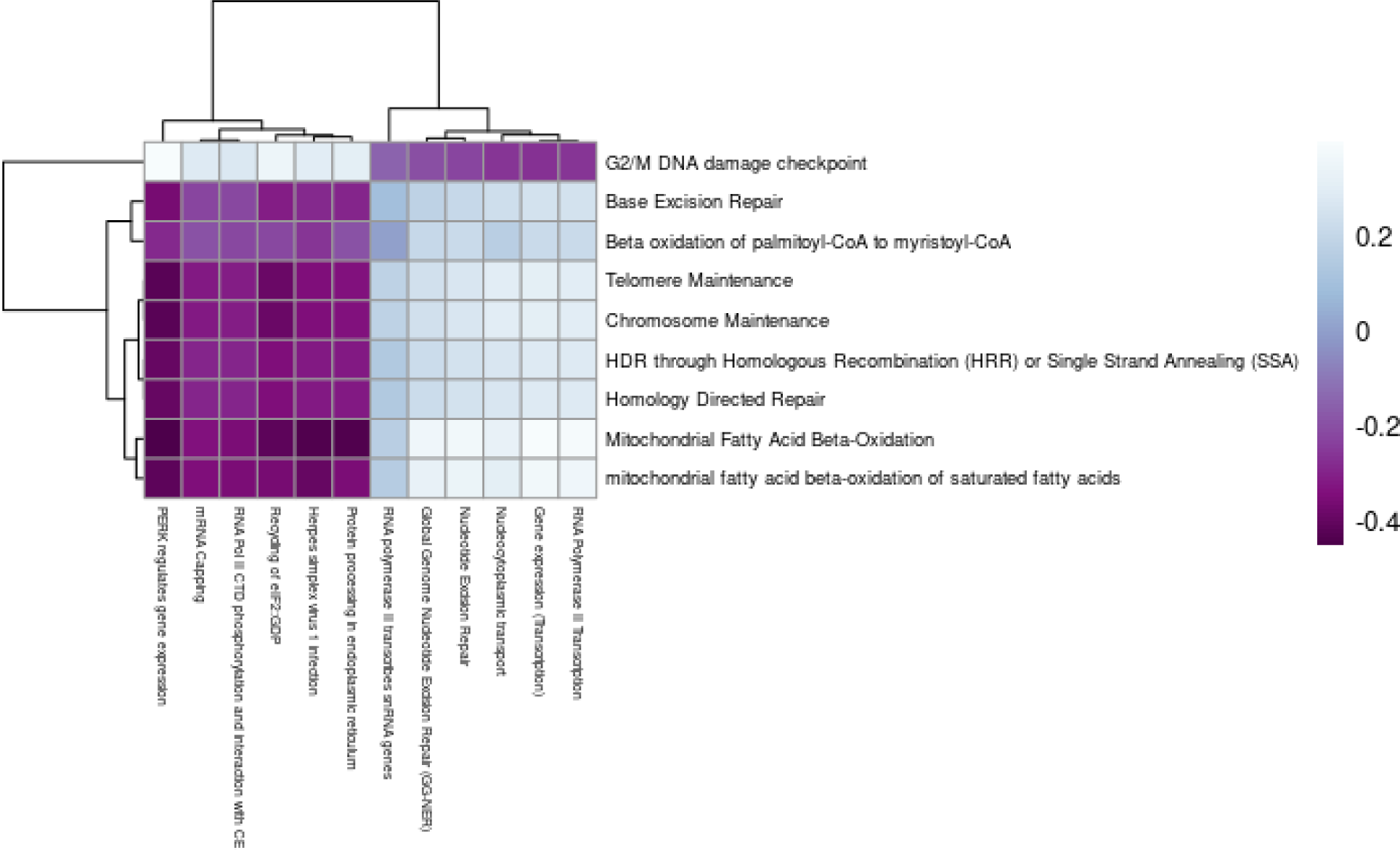
Correlations between pathway eigengenes in the hypothalamus (y-axis) and prefrontal cortex (x-axis). G2/M DNA damage checkpoint in the hypothalamus was negatively correlated with pathways related to ER stress and positively correlated with pathways related to RNA transcription in the prefrontal cortex. All other metabolic pathways in the hypothalamus were positively correlated with pathways related to ER stress and negatively correlated with pathways related to RNA transcription in the prefrontal cortex.

### 3.9 Data Resource

While we highlighted results from six tissues that stood out based on our filtering strategy, the data resource that accompanies this paper contains WGCNA and other analyses for each of the nine tissues, all the tissues combined, the brain tissues combined, and spleen, skeletal muscle, and small intestine combined. Additionally, analyses of all modules are present for the investigation of genes and networks not discussed in this paper. To ensure transparency and reproducibility a data resource was created to contain the analyses performed along with code, plots, and analysis description. The data resource can be intuitively navigated to meet the specific needs of individual researchers and is accessible through the link to the website (https://storage.googleapis.com/bl6_2dg_rnaseq/index.html).

The website is organized in a user-friendly manner with each WGCNA analysis housed under its own tab, classified by tissue or tissue combination. In addition to the WGCNA analysis, the website includes pages dedicated to the distribution analysis, outlier assessment, and sample information, as well as additional analyses performed for the paper. Researchers can access the raw code repository via a link to figshare containing the entire project with data, including the Shiny app, and a Shiny app for basic plots for any gene identified in this study.

## 4. Discussion

The systemic effects of 2DG across the body have applications for understanding the potential downstream effects of 2DG therapy, which has shown promise in treating cancer, epilepsy, polycystic kidney disease, autoimmune disease, and in delaying Alzheimer’s disease onset[1, 2, 4, 6, 7, 17, 47–50], and is currently being used to treat SARS-CoV-2[9, 11, 12]. Although 2DG therapy has shown promise in treating these conditions, there is lingering concerns regarding its safety. To better understand the potential risks associated with this treatment, our study investigated the systemic effects of 2DG on the transcriptomes of nine tissues in healthy young mice. By developing a novel filtering strategy of multiple statistical tests to identify robust genes and pathways affected by 2DG in multiple tissues, we found that WGCNA identified between 25 and 46 modules for each tissue, but only assessing modules with overrepresented pathways reduced the number of modules of interest for each tissue to between 12 and 32. Modules correlated with treatment further reduced the number of modules of interest to between zero and three for each tissue. Finally, ANOVA revealed that heart, hypothalamus, prefrontal cortex, small intestine, skeletal muscle, and liver were significantly associated with treatment and fit our filtering strategy. These six tissues were assessed for their robust 2DG effects, independently and collectively. Kidney, hippocampus, and spleen did not meet all the filtering criteria and therefore were not discussed (see data resource for their analysis).

2DG uniquely altered pathways and genes within each tissue module; however, tissue alterations could still be broadly identified as a result of glycolysis or N-linked glycosylation disruption. Glycolysis is necessary for energy production while N-linked glycosylation is necessary for protein function[39]. By analyzing the functions of our high confidence pathways, four tissues appeared to be affected by inhibition of glycolysis: heart, small intestine, hypothalamus, and liver. In contrast, we inferred that the remaining two tissues were affected by disruption of N-linked glycosylation: the prefrontal cortex and skeletal muscle.

Our study showed that 2DG treatment reduced the expression of gene pathways related to ATP production and glycolysis in the heart, both of which have been shown to play a role in heart failure[35, 51, 52]. Our results are consistent with a previous study[9], demonstrating that 2DG altered these pathways, but did not report any cardiac toxicity or significant changes in pathways related to ER stress. Although we did not perform EKGs, we observe no signs of heart trouble in the treated mice; moreover, patients with moderate to severe SARS-CoV-2 or advanced tumors treated with 2DG in clinical trials also showed no significant adverse heart-related effects due to this treatment[5, 10, 53]. Our findings suggest that there may be a strong association between ATP production and glycolysis pathways and the incidence of heart failure. Therefore, the reduction of these pathways by 2DG treatment may provide a new avenue for developing therapies to target these metabolic processes and prevent or treat heart failure.

While 2DG inhibits glycolysis, leading to a decrease in ATP production, it may also stimulate compensatory mechanisms that enhance oxidative phosphorylation, another pathway for ATP production[3, 9]. These findings suggest that 2DG primarily modifies metabolism in the heart and may have implications for developing new therapies for cardiovascular diseases, including using 2DG to improve heart health.

The small intestine is a unique immunological tissue because as it is the site in which immune cells, both sessile and recirculating, are continuously exposed to gut microfloral stimuli[54]. This encounter results in intestinal lymphocytes and leukocytes in an activated state that is not found in other organs of healthy mice. In agreement, we found that the small intestine was the only tissue in which untreated mice showed a very strong immune signature. Treatment with 2DG, in the small intestine resulted in a striking dampening of pathways associated with adaptive immunity and enrichment of innate immune pathways. Our analysis using CIBERSORT suggested that genes related to memory B cells and Th1 cells had decreased expression in mice treated with 2DG, which is in line with other studies showing a reduction in B and T cells and an increase in macrophages and eosinophils in the presence of 2DG[15, 55]. The increase in gene expression related to innate immunity pathways in the small intestine suggests that 2DG treatment may enhance the intestinal immune response against invading pathogens, while the decrease in gene expression related to adaptive immunity pathways may weaken the immune response against certain pathogens, potentially leading to increased susceptibility to infections. Additionally, our findings have implications for the potential relationship between the immune system and microbiome of the small intestine as the reduction in expression of memory B cell genes in mice treated with 2DG may promote dysbiosis. We envision two possibilities to explain the effects of 2DG on intestinal immune signature: 1) 2DG acts directly to eliminate on activated adaptive lymphocytes or their antigen presenting cells; and 2), 2DG eliminates gut bacteria that are the cause of immune stimulation. Our results emphasize the modulatory effects of 2DG on immunity in the small intestine, which may have implications for the development of new therapies for immune-related disorders.

Fatty acid oxidation is a critical process that helps to maintain energy homeostasis by providing energy substrates for ATP production[56]. The increase in fatty acid oxidation pathways in the hypothalamus with 2DG treatment may play a crucial role in the regulation of energy balance and metabolism when glycolysis is limited[57]. Our findings suggest that an increase in fatty acid oxidation pathways in the hypothalamus with 2DG may have positive implications for the overall health of the mice and inform new therapeutic approaches for metabolic disorders, such as obesity and diabetes, which are characterized by metabolic dysregulation[57]. This therapeutic approach may result in a shift towards energy expenditure and may help to counteract the effects of a high-fat diet or other metabolic disorders. However, further studies are needed to investigate the long-term effects of 2DG treatment on metabolic disorders in humans. Additionally, it is reported that treatment with 2DG in the 3xTgAD mouse model of Alzheimer’s disease increased expression of genes involved in ketone formation in the brain, reduced pathology, and resulted in increased serum ketone bodies[17]. Our study has shown that the hypothalamus switched to fatty acid oxidation in the presence of 2DG, suggesting a propensity of this critical anatomical structure to compensate for a reduction in glycolysis by utilizing fatty acids. Future studies could explore the potential benefits and limitations of targeting fatty acid oxidation pathways in the hypothalamus for the treatment of metabolic and neurodegenerative disorders.

Treatment with 2DG alters genes involved in protein metabolism in both the prefrontal cortex and skeletal muscle, suggesting potential implications for various diseases. In the prefrontal cortex, 2DG reduces gene expression in protein metabolism pathways in the ER compared to controls, which is associated with brain function regulation[58]. Previous studies have shown that by shutting down protein synthesis, 2DG can induce the unfolded protein response, reducing ER stress[14]. This is of particular interest as the prefrontal cortex is susceptible to dysregulation due to ER stress[59, 60], which is linked to neurological disorders such as Alzheimer’s and Parkinson’s disease[61]. Further investigations into the potential applications of altered protein metabolism pathways in the prefrontal cortex with 2DG treatment could be crucial for developing effective therapies for neurological disorders. Similarly, in skeletal muscle, prolonged ERK activation events were reduced in mice treated with 2DG compared to controls, which may have implications for metabolic health. Although ERK1/2 activation is necessary for maintenance of myofibers and neuromuscular synapses, prolonged activation can lead to muscle weakness[62–64]. Moreover, ERK pathway overactivation in skeletal muscle can lead to inflammation through activation of TNF-α, IL-6, IL-1ß, and contribute to metabolic disorders such as type 2 diabetes and obesity[65, 66]. This agrees with another study, which reported that 2DG decreases ERK pathways, resulting in decreased inflammatory markers[65]. Targeting ERK activation with 2DG in humans may have significant implications for the prevention and treatment of various diseases. Furthermore, the reduction of prolonged ERK activation events in the muscle and protein synthesis in the prefrontal cortex of mice treated with 2DG may also reduce inflammation and protect against misfolded proteins, which could be important for developing therapies for related diseases.

Our comparison between tissues showed that each tissue module was unique at the gene and pathway level demonstrating that 2DG affected each tissue in a unique manner. Although tissues responded uniquely, we did note potential relationships across tissues in response to 2DG. These findings demonstrate that the complex effects of 2DG on different tissues and pathways in mice, suggesting that further research is needed to understand the implications of these effects for overall health. However, our results also suggest that 2DG therapy has promising potential for the treatment of a range of diseases that affect different tissues and pathways in humans.

Although the present analysis sheds light on the effects of 2DG in male mice, there are several avenues for further investigation to expand our understanding of its potential benefits. First, it would be informative to explore whether female mice respond differently to 2DG. Second, due to the change in diet composition the effects of dosage length could not be assessed systemically. Third, while our filtering strategy identified robust tissues and module responses to 2DG, it is possible that other effects were not captured. Additionally, future research in disease models and in human patients is needed to fully understand the potential implications for clinical applications, and it should be noted that our study did not directly evaluate the safety of 2DG. Nonetheless, our findings improve our understanding of 2DG’s effects on healthy tissues, which may have implications for its safety in the long term.

## 5. Conclusion

Given the increasing interest in therapeutic use of 2DG, the goal of this study was to provide a systematic analysis of the transcriptomic consequences of glycolysis blockade by 2DG in major organs and tissues of healthy mammals. As we found surprisingly little transcriptional overlap among tissue types affected by 2DG treatment, the results are best explained by 2DG uniquely affecting each tissue type. The results are most consistent with two mechanisms through which 2DG is considered to act: inhibition of glycolysis and of N-linked glycosylation. This study provides a comprehensive resource on the effects of 2DG, a glycolytic inhibitor, on various tissues in non-diseased C57BL/6J mice, including a complete dataset, code, plots, and analysis descriptions for transparency and reproducibility. The website (https://storage.googleapis.com/bl6_2dg_rnaseq/index.html) is user-friendly, enabling easy navigation through all available analyses from all nine tissues. This system-wide characterization of 2DG activity across multiple tissues has the potential to inform its targeted therapeutic use, taking into account the mechanism relevant to each disease-affected tissue. However, further studies are needed to fully understand the mechanisms underlying the responses and determine their physiological significance.

**Funding:** This work was supported by NIH grant GM115518 to GWC., The Jackson Laboratory Director’s Innovation Fund (JAX-DIF 19000-18-19) and the Longevity Impetus Grant from Norn Group to C-HC.

### CRediT authorship contribution statement

**AEW:** Methodology, Writing-Original draft preparation, Software, Visualization, Formal analysis, Data Curation; **JJW:** Conceptualization, Methodology, Writing-Review and Editing, Project administration; **SEH:** Investigation, Writing-Review, and Editing; **JDS:** Investigation; **JW:** Investigation; **RP:** Conceptualization; **MWC:** Conceptualization, Writing-Review and Editing; **CCK:** Conceptualization, Methodology, Supervision, Resources, Funding acquisition; **DCR:** Conceptualization, Methodology, Resources, Writing-Review and Editing; **C-HC:** Conceptualization, Methodology, Supervision, Resources, Writing-Review and Editing, Project administration, Funding acquisition; **GWC:** Supervision, Resources, Writing-Review and Editing, Funding acquisition.

### Declaration of Competing Interest

The authors declare no conflict of interest.

## Abbreviations

2DG: 2-deoxy-D-glucose
RA: rheumatoid arthritis
SARS-CoV-2: severe acute respiratory syndrome coronavirus-2
SLE: systemic lupus erythematosus
ER: endoplasmic reticulum
WGCNA: weighted gene co-network analysis
ANOVA: analysis of variance
PCA: Principal component analysis
GSVA: Gene Set Variation Analysis
BCAA: branched-chain amino acids
NAD+: nicotinamide adenine dinucleotide

## Acknowledgements

We would like to thank Selcan Aydin for her invaluable discussions regarding the statistical approach and in proofreading the data resource. We thank Tamar Abel for proofreading the data resource. We thank Grace Stafford for aligning the transcriptomes and providing the processed RNA-seq data.

## Notes

https://storage.googleapis.com/bl6_2dg_rnaseq/index.html

### Competing Interest Statement

The authors have declared no competing interest.

### Summary of Updates

This version of the manuscript has been updated to include an additional author.

