## Supplemental figures for "Transcriptome Analysis Reveals Organ-Specific Effects of 2-Deoxyglucose Treatment in Healthy Mice"

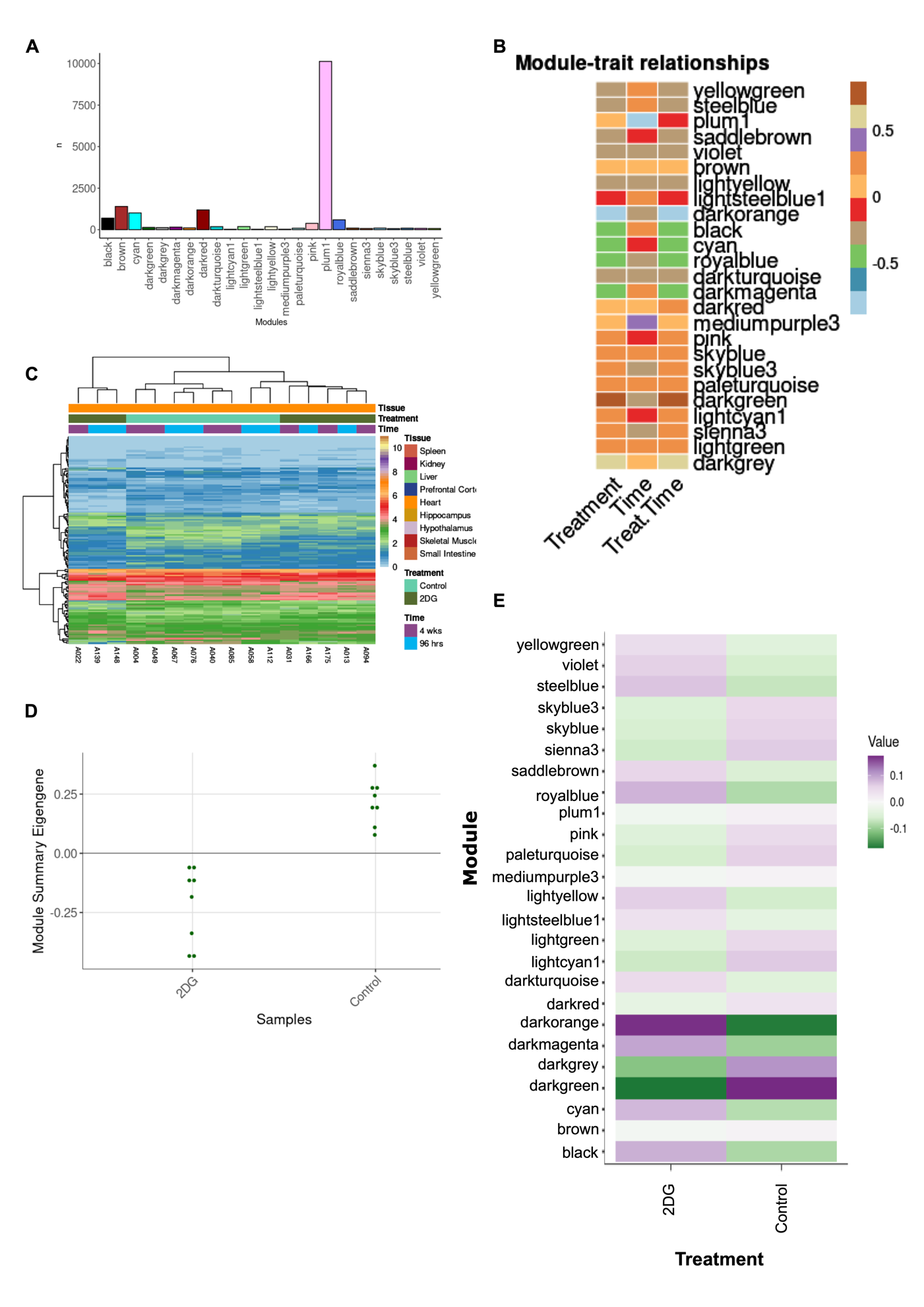

**Supplemental Figure 1:** The heart darkgreen module showed: A) A total of 25 modules were identified, each containing between 27 and 10,127 genes. B) Correlation analysis between time, treatment, and treatment in a time dependent manner revealed three modules significantly correlated with treatment. C) The average eigengene expression of the darkgreen module shows that expression levels are decreased in mice treated with 2DG compared to control mice. D) Summary eigengene expression for each sample in the darkgreen module confirms decreased expression in mice treated with 2DG compared to control mice. E) Individual genes across samples clustered by treatment with two clusters containing mice treated with 2DG.

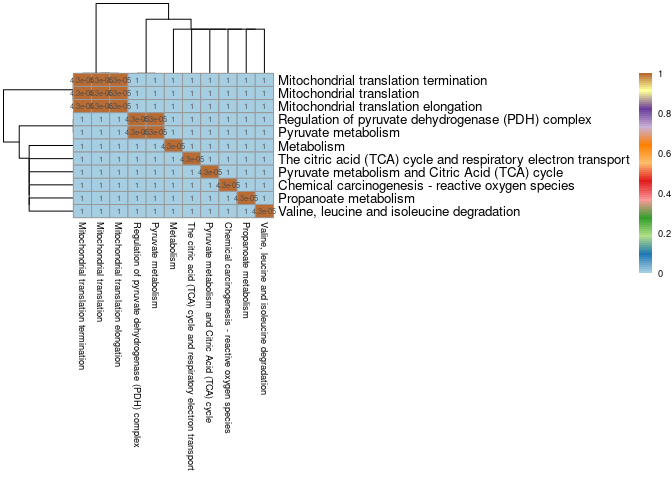

**Supplemental Figure 2:** Comparison of genes identified within each pathway in the heart darkgreen module show that each pathway contains a unique set of genes. Pathways are generally comprised of distinct genes, as depicted by a Jaccard similarity matrix. Genes present within each overrepresented pathway are compared. Brown indicates significant overlap and blue indicates non-significant overlap between two sets of genes.

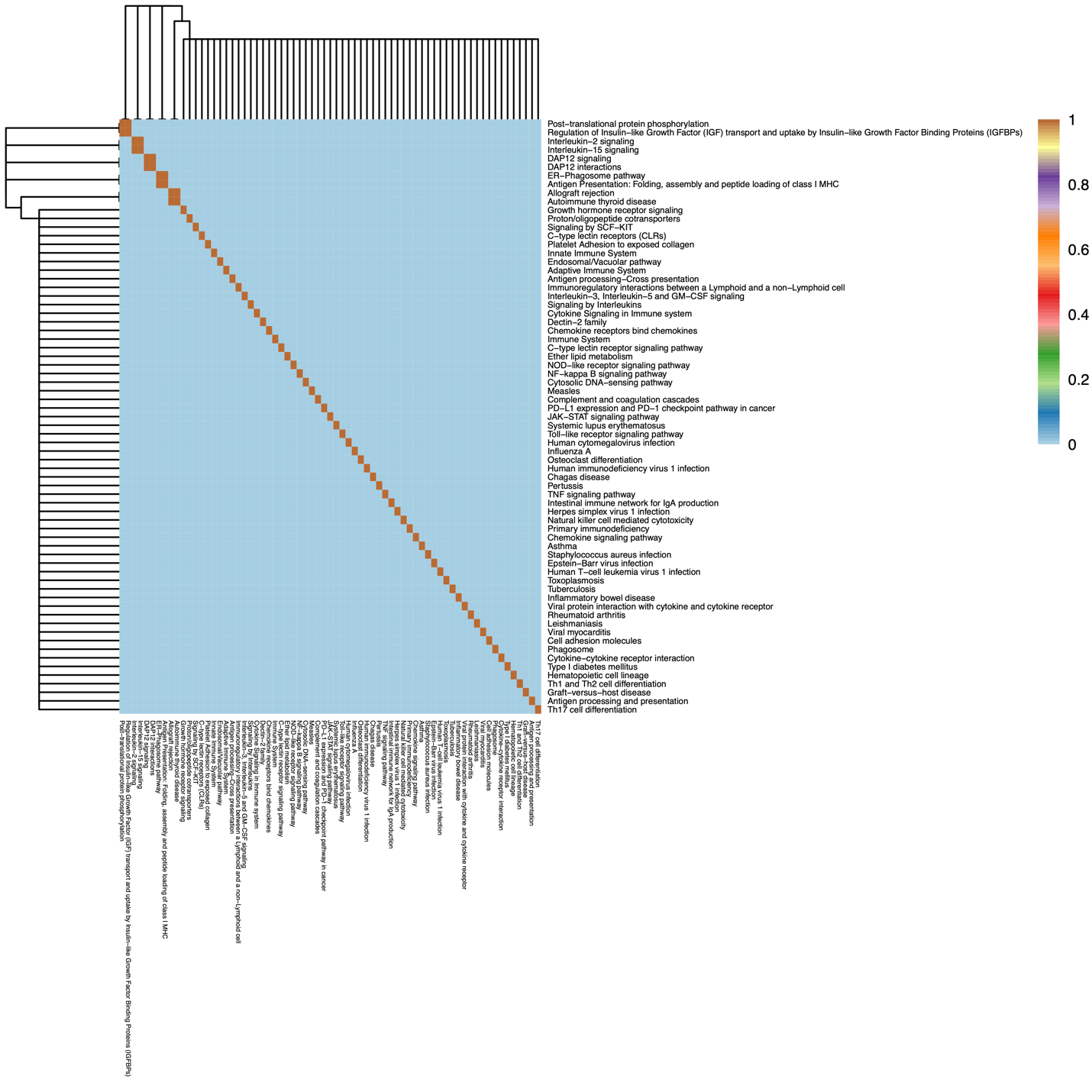

**Supplemental Figure 3:** Comparison of genes identified within each pathway in the small intestine green4 module show that each pathway contains a unique set of genes. Pathways are generally comprised of distinct genes, as depicted by a Jaccard similarity matrix. Genes present within each overrepresented pathway are compared. Brown indicates significant overlap and blue indicates non-significant overlap between two sets of genes.

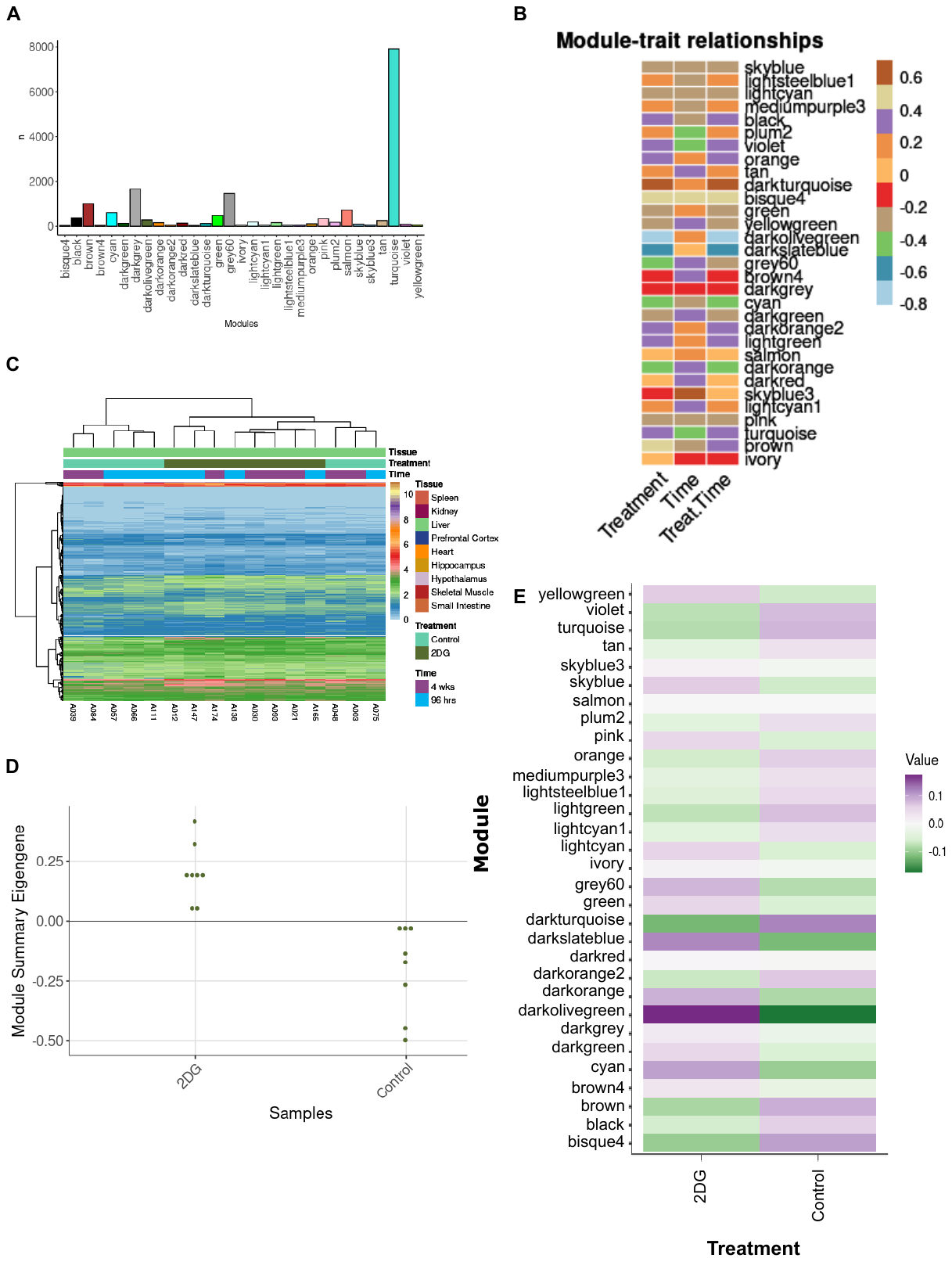

**Supplemental Figure 4:** The darkolivegreen module in the liver showed: A) A total of 31 modules were identified in the liver, each containing between 41 and 7,909 genes. B) Correlation analysis between time, treatment, and treatment in a time dependent manner revealed three modules significantly correlated with treatment. C) The average eigengene expression of the darkolivegreen module shows that expression levels are increased in mice treated with 2DG compared to control mice. D) Summary eigengene expression for each sample in the darkolivegreen module confirms increased expression in mice treated with 2DG compared to control mice. E) Individual genes across samples clustered by treatment except for three control sample, which clustered with 2DG-treated mice.

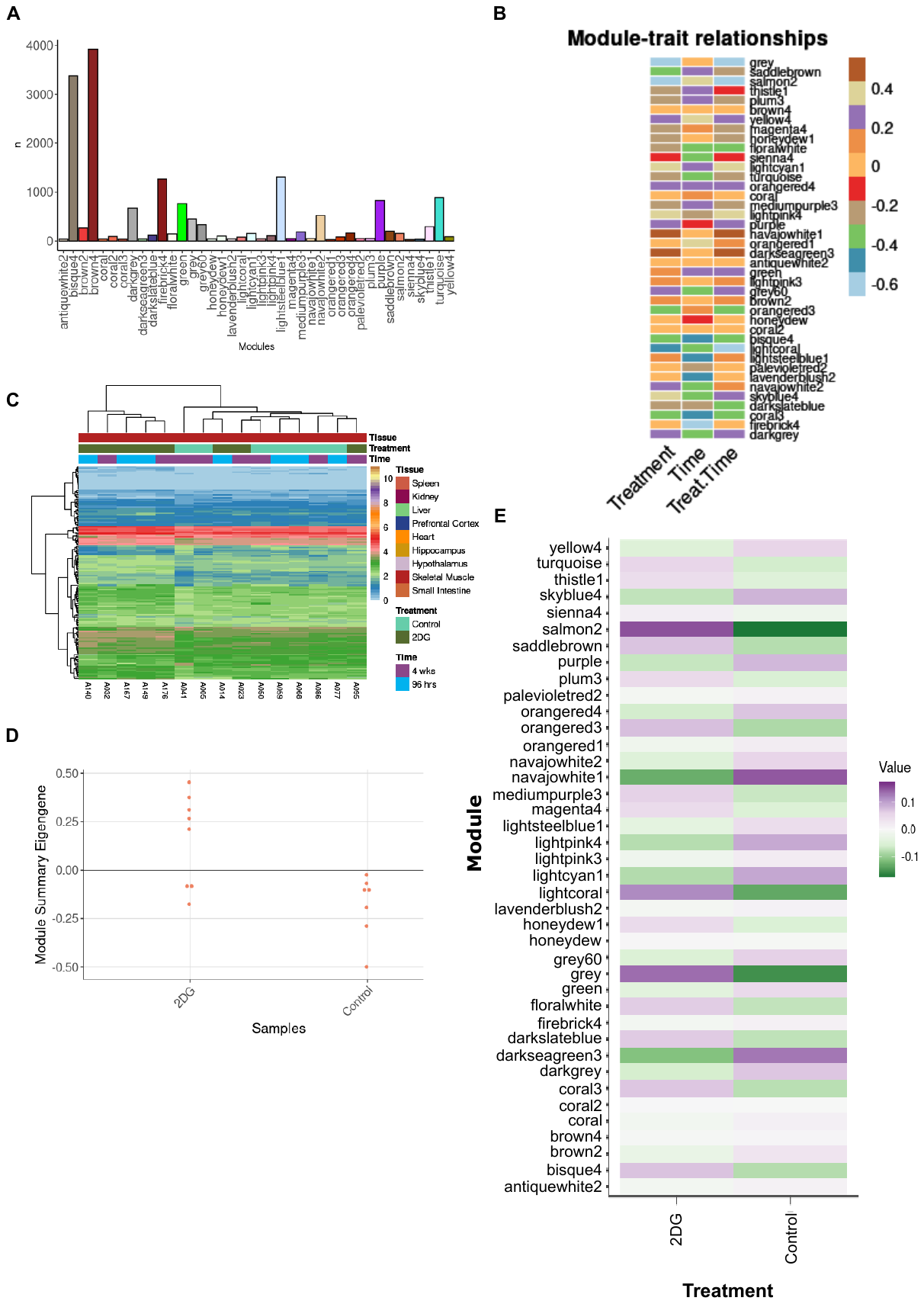

**Supplemental Figure 5:** The salmon2 module in skeletal muscle showed: A) A total of 40 modules were identified in the skeletal muscle, each containing between 28 and 3,920 genes. B) Correlation analysis between time, treatment, and treatment in a time dependent manner revealed three modules significantly correlated with treatment. C) The average eigengene expression of the salmon2 module shows that expression levels are increased in mice treated with 2DG compared to control mice. D) Summary eigengene expression for each sample in the salmon2 module confirms increased expression in mice treated with 2DG compared to control mice. E) Individual genes across samples clustered by treatment except for three 2DG-treated mice, which clustered with control mice.

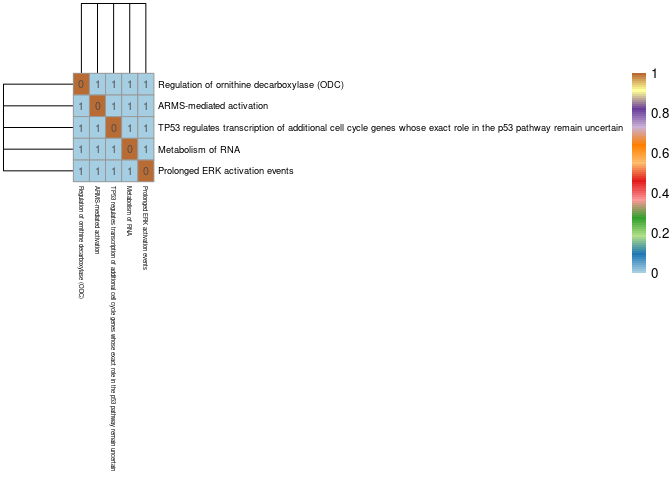

**Supplemental Figure 6:** Comparison of genes identified within each pathway in the skeletal muscle salmon2 module show that each pathway contains a unique set of genes. Pathways are generally comprised of distinct genes, as depicted by a Jaccard similarity matrix. Genes present within each overrepresented pathway are compared. Brown indicates significant overlap and blue indicates non-significant overlap between two sets of genes.

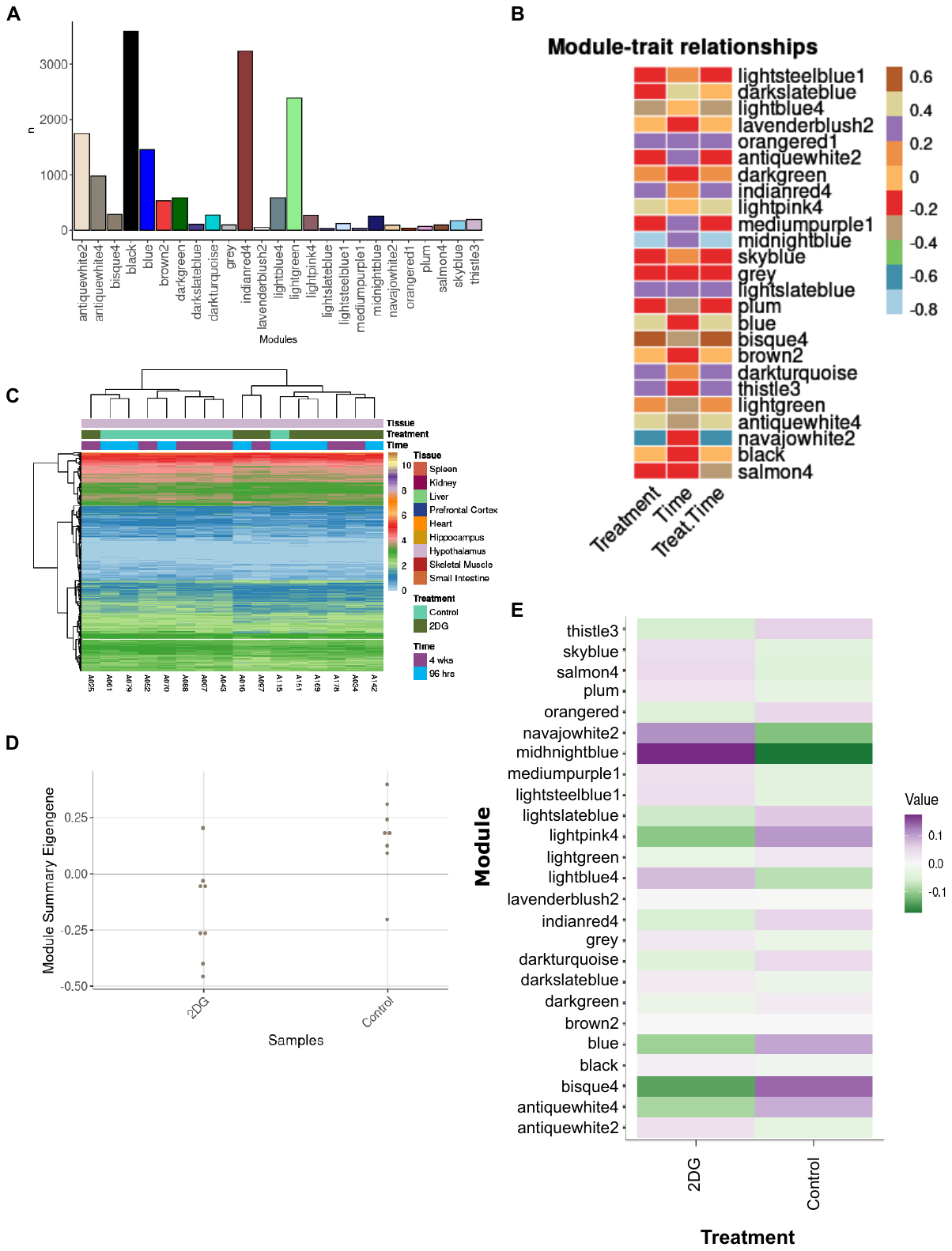

**Supplemental Figure 7:** The bisque4 module in the hypothalamus showed: A) A total of 25 modules were identified in the hypothalamus, each containing between 27 and 3,598 genes. B) Correlation analysis between time, treatment, and treatment in a time dependent manner revealed four modules significantly correlated with treatment. C) The average eigengene expression of the bisque4 module shows that expression levels are decreased in mice treated with 2DG compared to control mice. D) Summary eigengene expression for each sample in the bisque4 module confirms decreased expression in mice treated with 2DG compared to control mice. E) Individual genes across samples clustered by treatment with the exception of two samples one control and one 2DG sample, each clustered with the opposite mouse group.

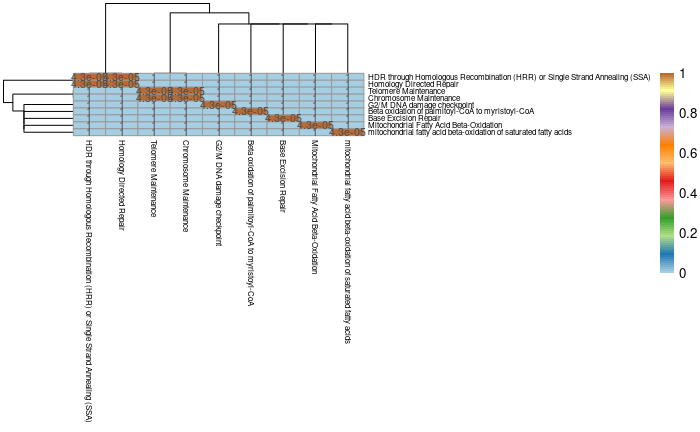

**Supplemental Figure 8:** Comparison of genes identified within each pathway in the hypothalamus bisque2 module show that each pathway contains a unique set of genes. Pathways are generally comprised of distinct genes, as depicted by a Jaccard similarity matrix. Genes present within each overrepresented pathway are compared. Brown indicates significant overlap and blue indicates non-significant overlap between two sets of genes.

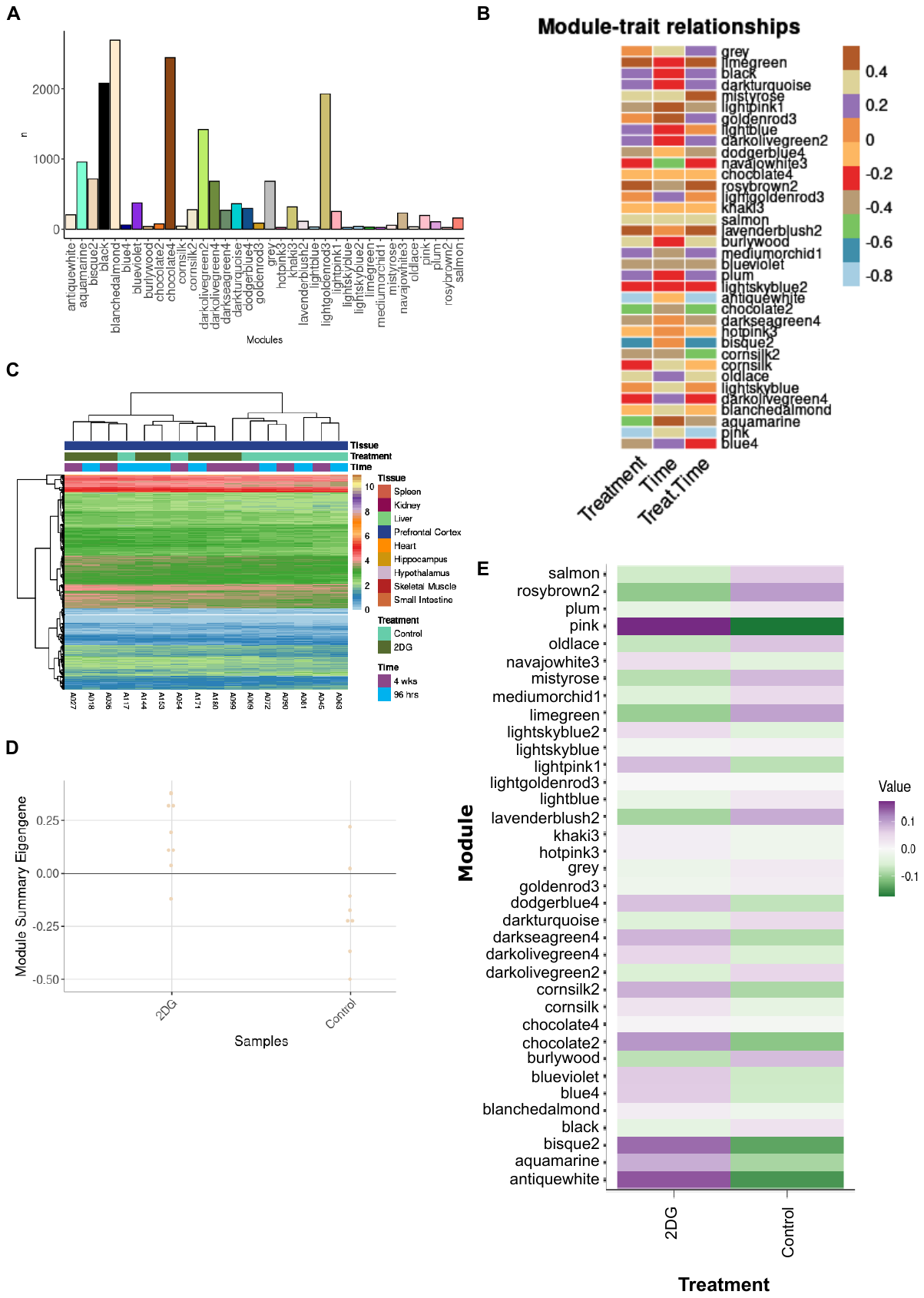

**Supplemental Figure 9:** The bisque2 module in the prefrontal cortex showed: A) A total of 36 modules were identified in the prefrontal cortex, each containing between 22 and 2,698 genes. B) Correlation analysis between time, treatment, and treatment in a time dependent manner revealed five modules significantly correlated with treatment. C) The average eigengene expression of the bisque2 module shows that expression levels are increased in mice treated with 2DG compared to control mice. D) Summary eigengene expression for each sample in the bisque2 module confirms increased expression in mice treated with 2DG compared to control mice. E) Individual genes across samples clustered by treatment with the exception of three samples, two control mice clustered with 2DG-treated mice and one 2DG-treated mice clustered with control mice.

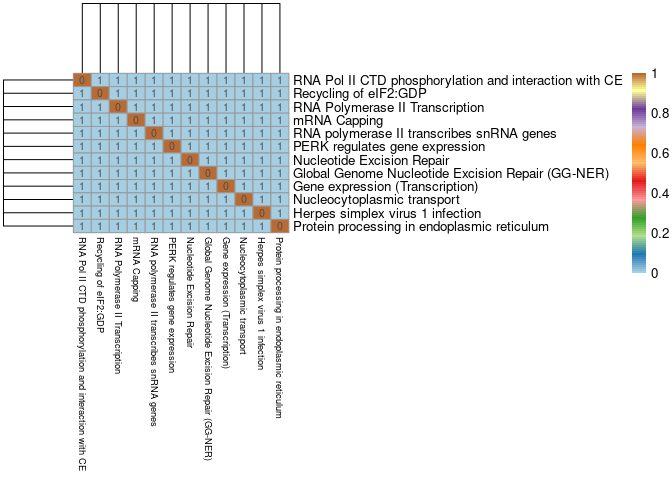

**Supplemental Figure 10:** Comparison of genes identified within each pathway in the prefrontal cortex bisque4 module show that each pathway contains a unique set of genes. Pathways are generally comprised of distinct genes, as depicted by a Jaccard similarity matrix. Genes present within each overrepresented pathway are compared. Brown indicates significant overlap and blue indicates non-significant overlap between two sets of genes.

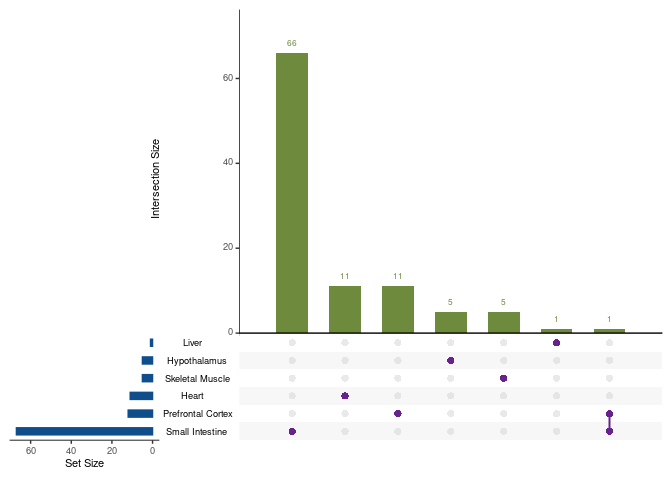

**Supplemental Figure 11:** Number of overrepresented pathways shared between tissue modules. Each dot represents the tissue or tissues being compared. There were very few pathways shared across tissues and no pathway was shared across all tissues. Only tissue combinations with one or more shared genes are shown.
